# Hepatic stellate cells drive hypoxia-inducible factor (HIF)-dependent metabolic rewiring and proliferation in hepatocellular carcinoma cells

**DOI:** 10.64898/2026.10.26.761674

**Authors:** Demi J. Rijlaarsdam, Jung-Chin Chang, J. Bernd Helms, Esther A. Zaal, Celia R. Berkers

## Abstract

Hepatocellular carcinoma (HCC) is the most prevalent primary liver cancer in adults and represents the third most common cause of cancer-related deaths worldwide. With limited treatment options, HCC remains a major global health problem. The majority of HCC develops in a cirrhotic liver, where the tumor microenvironment (TME) is enriched in fibroblasts derived from activated hepatic stellate cells (HSCs). Cell-cell interactions in TME play an important role in metabolic reprogramming, which is increasingly recognized as a crucial driver in HCC proliferation. The tumor-promoting and metabolic effects of HSCs on HCC cells however remain incompletely characterized. Understanding how HSC-HCC interaction regulates HCC proliferation can lead to identification of novel therapeutic targets. Here, we established a Transwell co-culture model of HSCs and HCC cells to investigate how HSC-HCC interactions regulate HCC metabolism and proliferation. Using isotope tracing metabolomics, we show that co-culture with HSCs increases proliferation of HCC cells by rewiring glucose metabolism to increase glycolysis and *de novo* synthesis of nucleotides. Transcriptomic and immunofluorescence staining analyses revealed that HSCs upregulate hypoxia-inducible factor (HIF) signaling in HCC cells, which contributes to metabolic rewiring and HCC proliferation. We further identified H_2_O_2_ as a candidate crosstalk signal that could mediate HSC-dependent activation of HIF signaling in HCC cells. Together, these findings reveal a previously overlooked modulatory role of HSC-HCC interaction in metabolism-driven HCC progression. Targeting these pathways may represent a promising strategy to improve HCC treatment.

## Introduction

Liver cancer represents the third most common cause of cancer-related deaths worldwide and is the sixth most commonly diagnosed cancer, with hepatocellular carcinoma (HCC) accounting for more than 80% of liver cancers (Singal et al., 2023). HCC is a growing global health problem, with rising incidence and mortality rates over the past two decades and mortality rates predicted to reach 1 million deaths annually by 2030 (Kim & Viatour, 2020; Qiu et al., 2024). Early-stage HCC is often clinically asymptomatic, leading to delayed diagnosis and advanced disease at initial presentation. Its limited response to conventional chemotherapy and radiotherapy further contributes to the paucity of effective therapeutic options and the poor prognosis of patients with advanced disease (Kim & Viatour, 2020; Llovet et al., 2008, 2022; Mauro et al., 2025; Wang & Deng, 2023). When detected at an early stage, treatment options for HCC include surgical resection and liver transplantation, but for advanced-stage and recurrent HCC, therapy is limited to systemic treatments. Currently, the most effective systemic treatment for advanced HCC consists of atezolizumab, an immune checkpoint inhibitor, and bevacizumab, an anti-angiogenic monoclonal antibody. Patients however face severe adverse effects while long-term survival remains limited (Cerreto et al., 2023; Cheng et al., 2022; Finn et al., 2020; Manfredi et al., 2025; Selene et al., 2024). This illustrates the urgent need to identify novel targets and effective treatment strategies for HCC patients.

The dynamic crosstalk between cancer cells, immune cells and stromal cells in the HCC tumor microenvironment (TME) is of great importance to drive the malignant progression of HCC (Yu et al., 2022). HCC primarily emerges in severely fibrotic or cirrhotic livers (M. C. Wallace & Friedman, 2014). Hepatic stellate cells (HSCs), liver fibroblasts, are directly involved in shaping the microenvironment of these pre-malignant stages. Upon liver injuries, HSCs are activated and transdifferentiated into myofibroblasts which secrete various factors that ensure extracellular matrix (ECM) deposition (Molenaar et al., 2023; Tsuchida & Friedman, 2017). In addition, HSCs may play a role in HCC progression, where it recently has been shown they represent a major source of cancer-associated fibroblasts (CAFs) (Cogliati et al., 2023; Ying et al., 2023). CAFs in turn contribute to liver cancer maintenance and progression by directly promoting proliferation and angiogenesis, as well as by remodeling the TME through the deposition of collagen, one of the main components of the ECM (Cogliati et al., 2023). While the mechanisms through which HSCs drive HCC progression remain to be fully elucidated, HSC-secreted factors such as TGF-β, FGF and HGF, ECM-remodeling enzymes, and an upregulation and secretion of angiogenic factors can promote HCC progression within the TME (Han et al., 2014; Lin et al., 2020; Y.-N. Ma et al., 2021).

An altered metabolic state is considered a key hallmark of cancer. Metabolic rewiring is essential for providing energy, building blocks and redox homeostasis required for survival, rapid proliferation and drug resistance (Pavlova et al., 2022). Like many other tumor types, HCC is associated with extensive metabolic reprogramming, which has been linked to a poor prognosis. This reprogramming includes major metabolic pathways such as glucose, amino acid and lipid metabolism (Park & Hall, 2025; Satriano et al., 2019). As such, targeting cancer-associated metabolic alterations have emerged as a promising therapeutic strategy for HCC (An & Li, 2025; Du et al., 2022; Gnocchi et al., 2023). Stromal cells such as CAFs have been suggested to support cancer metabolism within the liver TME, for example by releasing glucose-derived metabolites such as pyruvate and lactate that can be used by the cancer cell (F.-Q. Xu et al., 2023). HSCs also secrete extracellular vesicles containing hexokinase 1 (HK1), an important glycolytic enzyme, which HCC cells can subsequently hijack to promote proliferation (Q. Chen et al., 2022). While these observations underscore the importance of HSC-HCC metabolic crosstalk within the TME, our understanding of these interactions and HSC-mediated metabolic rewiring in HCC remains limited, hindering the identification of metabolic targets that could be therapeutically exploited to improve current HCC treatment.

Here, we established a Transwell co-culture model to investigate HSC-HCC metabolic crosstalk and its functional and metabolic effects on HCC cells. We show that HSCs induce a hypoxia-inducible factor (HIF)-mediated metabolic rewiring in glucose metabolism in HCC, resulting in increased proliferation. We suggest a key role of HSC-derived reactive oxygen species (ROS) in mediating the intercellular communication between HSCs and HCC cells. These findings reveal an HSC-driven metabolic adaptation in HCC cells that promotes cancer progression and may provide new strategies for therapeutic interventions.

## Materials and methods

### Cell lines and culture conditions

The mouse HCC cell line Hepa1-6 was a kind gift from Tytgat Institute for Liver and Intestinal Research, Amsterdam UMC. The mouse HSCs (mHSCs) cell line was established from quiescent primary mHSCs. Briefly, quiescent primary mHSCs were isolated by 3-step EGTA-pronase-collagenase liver perfusion and Nycodenz gradient as previously described (Mederacke et al., 2015). The purity of quiescent primary mHSCs were verified by large UV-positive lipid droplets. The quiescent primary mHSCs were activated by culturing on cell culture-treated hard plastic surface in DMEM with 10% fetal bovine serum (FBS, Thermo Fisher Scientific, #A5256701) and penicillin–streptomycin. The mHSCs cell line was subsequently established by self-immortalization. Both cell lines were adapted to and maintained in William’s E Medium (Thermo Fisher Scientific, #A12176), supplemented with 10% FBS and 1X GlutaMAX (Thermo Fisher Scientific, #35050061). Cells were cultured at 37°C under 5% CO_2_, and tested every 2 months for mycoplasma contamination.

### Co-culture model

Costar® sterile and tissue culture treated 24-well cell culture plates and 6.5 mm Transwell® inserts with a permeable polyester membrane and a 0.4 µm pore size (Corning, #3470) were used. One day prior to the start of the co-culture, Hepa1-6 cells were seeded in the basolateral compartment (bottom well) at a density of 20,000 cells per well in 750 µL medium. In parallel, mHSCs were seeded in the apical compartment (insert) in a separate plate at a density of 100,000 cells per insert to achieve 100% confluency, in 150 µL medium with 600 µL added to corresponding bottom wells. At the start of co- and mono-culture, inserts were placed in the wells with Hepa1-6 cells, while culture media was refreshed with 600 µL in the basolateral compartment and 150 µL in the apical compartment. For mono-culture conditions, the same procedures were followed, with no cells in either insert or bottom well. Cells were co- and mono-cultured for 72 hours in triplicate wells per condition.

### Cell proliferation assay

To measure proliferation of both cell types in the co-culture model, inserts were carefully separated from bottom wells. Cells were washed with 1X phosphate-buffered saline (PBS) and detached using 0.05% Trypsin-EDTA (1X) (Thermo Fisher Scientific, #25300), after which cell culture medium was added to neutralize trypsin. Cells were collected for cell counting using a hemocytometer, from which total cell numbers per well and fold changes relative to seeding densities of co- and mono-cultured cells were calculated.

In addition, proliferation was assessed of co- and mono-cultured cells treated with the following inhibitors: glycolysis inhibitor 2-deoxy-D-glucose (2-DG) (Sigma-Aldrich, #D6134), glucose transporter 1 (GLUT1) inhibitor BAY-876 (MedChemExpress, #HY- 100017), HIF-1α suppressing agent PX-478 (MedChemExpress, #HY-10231) and HIF-2α inhibitor PT-2385 (MedChemExpress, #HY-12867). A combination of 75 µM ascorbate (Sigma-Aldrich) and 500 µM 2-phosphate-L-ascorbate (Sigma-Aldrich), to keep intracellular vitamin C levels stable for 72 hours of culture (Frikke-Schmidt & Lykkesfeldt, 2010), was applied for inhibition of oxidative stress and HIF signaling. These inhibitors were administered in concentrations determined by prior titration and viability analysis in both Hepa1-6 cells and mHSCs. At the start of the co- and mono-cultures, treatments were started including a vehicle control. After the 72 hours of culturing according to our standard co-culture model procedures, proliferation rates were determined.

### Transcriptomics

Samples for RNA sequencing were harvested from cells in the co-culture model by separating inserts from bottom wells. RNA harvesting and isolation was done using the RNeasy Micro Kit (Qiagen, #74004), according to the protocol of the manufacturer. Next generation sequencing was performed by the Utrecht Sequencing Facility (USEQ): RNA quality was checked with the Agilent Fragment Analyzer 5300 system using the RNA Kit (15 nt) (Cat. DNF-471-1000). RNA quantity was measured with the Invitrogen™ Qubit™ Fluorometer using the Qubit RNA HS Assay Kit (Cat. Q32855). 100 ng of total RNA was used to prepare TruSeq Stranded mRNA libraries (Cat. 20020594) following the manufacturers protocol, using half the reagents volume and with custom 384 xGen UDI-UMI adapters from IDT. After the library preparation, libraries were checked with the Fragment Analyzer system dsDNA 910 Reagent Kit (35-1500 bp) (Cat. DNF-910-K1000) and with Qubit dsDNA HS Assay Kit (Cat. Q32854). Sample libraries were pooled equimolar. Libraries were sequenced on a Nextseq2000 (Illumina) by using a P2 flowcell with 50 bp paired-end reads. Quality control on the sequence reads from the raw FASTQ files was done with FastQC (v0.11.9).

TrimGalore (v0.6.7) was used to trim reads based on quality and adapter presence after which FastQC was again used to check the resulting quality. rRNA reads were filtered out using SortMeRNA (v4.3.6) after which the resulting reads were aligned to the reference genome fasta (Mm_GRCm38_gatk_sorted.fasta) using the STAR (v2.7.10b) aligner. Follow-up QC on the mapped (bam) files was done using Sambamba (v0.8.2), RSeQC (v5.0.1) and PreSeq (v3.2.0). Readcounts were then generated using the Subread FeatureCounts module (v2.0.3) with the Mus_musculus.GRCm38.70.gtf gtf file as annotation, after which normalization was done using the R-package edgeR (v3.40). Differential Expression analysis was performed with an inhouse R-script using DESeq2 (v1.38.0), taking the raw readcounts as input. Finally a summary report was created using MultiQC (v1.14). Subsequent analyses were performed using the Utrecht University ATWiCS ShinyApp including normalization of the raw readcounts. Here, a differential expression (DE) analysis was performed, and the genes that were significantly differential between conditions with a log2-transformed fold change of at least (-)0.25 were used for pathway analyses in Enrichr (E. Y. Chen et al., 2013; Kuleshov et al., 2016; Xie et al., 2021).

### Liquid chromatography-mass spectrometry (LC-MS)-based semi-targeted metabolomics

Stable isotope tracing was performed as described (Zaal et al., 2025). 24 hours prior to harvesting from co- and mono-cultures, media were replaced with culture medium (Williams’ Medium E, with sodium bicarbonate, without L-glutamine, glucose, sodium pyruvate, phenol red (USBiological, #W1105-02), supplemented with glutamine, sodium pyruvate and FBS as standard culturing conditions), and with [U-^13^C_6_]-D-glucose (Cambridge Isotope Laboratories, #CLM-1396). When testing inhibitors, this [U-^13^C_6_]-D-glucose medium was supplied at the start of co- and mono-cultures, at the same time of inhibitor administration (or vehicle control), 72 hours prior to harvesting. The GLUT1 inhibitor BAY-876 and HIF-2α inhibitor PT-2385 were used, as also employed in the cell proliferation assay. To harvest extracellular metabolites, medium samples were obtained from the basolateral compartment of the Transwell model for each condition. To extract the metabolites of all medium samples, 10 µL of the harvested medium was diluted in 1 mL of freshly prepared ice-cold MS lysis buffer (methanol/acetonitrile/ULC/MS-grade H_2_O, 2:2:1). To harvest intracellular metabolites, the cell types in co-culture were first separated from each other, while keeping cells cold to ensure no metabolic changes were induced upon handling of the cells. Before extraction, cells were washed with ice-cold PBS. Then, 150 µL of ice-cold MS lysis buffer was added to the cells. Subsequently, cells were kept for 15 minutes at 4°C while slowly rotating to extract all intracellular metabolites, which was collected in Eppendorf tubes. All samples were shaken for 10 minutes at 4°C and centrifuged at 14,000 x g for 15 minutes at 4°C, after which the supernatant was collected for LC-MS.

LC-MS analysis was performed on an Exactive mass spectrometer (Thermo Scientific) coupled to a Dionex Ultimate 3000 autosampler and pump (Thermo Scientific) or Q-Exactive HF mass spectrometer (Thermo Scientific) coupled to a Vanquish autosampler and pump (Thermo Scientific). The MS operated in polarity-switching mode with spray voltages of 4.5 kV and −3.5 kV. Metabolites were separated using a Sequant ZIC-pHILIC column (2.1 x 150 mm, 5 μm, guard column 2.1 x 20 mm, 5 μm; Merck) with elution buffers acetonitrile (A) and eluent B (20 mM (NH4)2CO3, 0.1% NH4OH in ULC/MS-grade water (Biosolve)). Gradient ran from 20% eluent B to 60% eluent B in 20 minutes, followed by a wash step at 80% and equilibration at 20%. Flow rate was set at 100 μL/min. Analysis was performed using Tracefinder software (Thermo Scientific). Metabolites were identified and quantified on the basis of exact mass within 5 ppm and further validated by concordance with retention times of standards. Peak intensities were normalized based on total ion count. In ^13^C-tracer experiments, isotopomer distributions were corrected for the natural abundance of ^13^C. Pathway enrichment analysis was performed using MetaboAnalyst 5.0/6.0 (Pang et al., 2021, 2024). Extracellular metabolite levels were used to calculate uptake or secretion, where Δ levels were calculated by subtracting the mean level in medium-only controls from each of the replicates.

### Immunofluorescence (IF) staining and nuclear signal quantification of HIF-1α

Cells were seeded on cover slips (⌀ 12 mm), while standard co-culture model procedures were followed as described above. After the 72 hours of co- and mono-culturing, cells were washed with 1X PBS and fixed with 4% (w/v) paraformaldehyde (PFA). This was incubated for 15 minutes at room temperature before decanting fixative and washing cells 3 times with 1X PBS. Six hours prior to fixing cells, freshly prepared 1 mM dimethyloxallyl glycine (DMOG, MedChemExpress, #HY-15893) was added to one condition of Hepa1-6 cells without co-culture to ensure HIF-1α stabilization and accumulation in Hepa1-6, to serve as a positive control. Staining was done by first permeabilizing the cells using 1% Triton X-100 in 1X PBS for 10 minutes at room temperature. Next, the permeabilization solution was discarded and cells were washed 3 times with a phosphate-buffered saline with 0.05% Tween 20 (PBST) solution (5 minutes of incubation per wash). In-well staining was performed, using the primary HIF-1α polyclonal antibody (Novus Biologicals, #NB100-449) in 1:100 dilution in PBST. This was incubated overnight at 4°C and cells were washed 3 times with PBST. Then, the secondary antibody (goat-anti-rabbit, Alexa488) was added in 1:100 dilution in PBST. Phalloidin was also added in the secondary antibody incubation step (1:100 in PBST, Alexa Fluor 568), to counterstain the cell body around the nucleus (F-actin). Secondary only controls were included for all analyses, and all were negative for signal. While protected from light in the following procedures, plates were shaken at room temperature for 1 hour. Coverslips were then washed 3 times with PBST, 5 minutes of incubation per wash, before rinsing cells once with Milli-Q. After blotting away excess liquid, coverslips were mounted on a slide using VECTASHIELD Antifade Mounting Medium With DAPI (Brunschwig Chemie, #H-1200), fluorescently staining the nucleus of cells. The slides are stored at 4°C and protected from light before subsequent microscopy analyses.

Images were collected under consistent settings with a NIKON A1R confocal microscope using a 40x Plan Apochromat air objective (NA 0.95) with a laser set of 405, 488, 561 and 645 nm. A quad band dichroic (405/488/561/640 nm) and emission filters were positioned to detect, Alexa568 (595/30) and DAPI (450/50) simultaneously and Alexa488 (515/30) in a sequential line mode. The pinhole was set to 38.31 um (∼1.5 AU), laser powers were 3.7, 1.1 and 1.4% respectively and the pixel size 310 nm. The system was run by NIS elements software (NIKON instruments, version 5.1).

ImageJ was used to process images, where a DAPI-derived nuclear mask was created and applied to the Alexa488 signal to present nuclear HIF-1α. Nuclear HIF-1α intensities were calculated as follows: single cell intensity ratios were calculated using Cellprofiler (Kamentsky et al., 2011) (version 4.2.8) using the pipeline supplied in their supplementary data. Briefly, images were preprocessed by smoothing, intensity rescale and correction of illumination shading. Nuclei were segmented by Otsu‘s thresholding and splitting was performed based on the background levels. Resulting objects were filtered to have an area above 250 pixels. These objects were the seeds to define cell outlines in the combined actin and DAPI channels, cells with aberrant nuclear shapes were excluded and filtered by area as well (>2000 pixels). Finally, the mean nuclear intensities of Alexa488 labeled HIF-1α were measured and analyzed.

### Detection of H_2_O_2_ secretion by homovallinic acid (HVA)-horseradish peroxidase (HRP) method

Accumulative secretion of H_2_O_2_ into media by Hepa1-6 cells and mHSC in mono- and co-culture was detected by the formation of fluorescent HVA dimer from HVA (Sigma, #H1252) and H_2_O_2_ in the presence of HRP (Sigma, #10108090001) as previously described with modification (Ruch et al., 1983). Mono- and co-cultures of Hepa1-6 and mHSC in Transwells were set up as described above. The assay was performed in a modified Hank’s buffered salt solution for 5% CO_2_ (107.8 mM NaCl, 21.43 mM NaHCO_3_, 4.56 mM KCl, 1.27 mM CaCl_2_, 0.41 mM MgSO_4_, 0.39 mM MgCl_2_, 0.45 mM NaH_2_PO_4_, 0.34 mM Na_2_HPO_4_, 10 mM HEPES-NaOH [pH 7.4]) pre-equilibrated in a 5% CO_2_ incubator before use (Chang et al., 2016). After removing the spent media, the inserts and wells were washed twice with the experimental medium to remove serum. Eight hundred microliter and 200 µL of 1X HVA-HRP solution (100 µM HVA and 1 U/mL HRP in the experimental medium) was added to the inserts and wells respectively and 100 µL medium was harvested from the well as baseline. After 18 hours, 100 µL of medium was harvested from the wells as well as from the inserts. All harvested medium was immediately mixed with 10 µL 250 mM EDTA-NaOH (pH 12.0) to inactivate HRP and maximize fluorescence of HVA homodimer. An H_2_O_2_ standard curve (final concentrations 10 µM to 0.3125 µM; freshly prepared from 30% [w/w] H_2_O_2_ solution) was constructed in 1X HVA-HRP solution to quantify medium H_2_O_2_. To compare H_2_O_2_ secretion in Hepa1-6 and mHSC mono-cultures, the accumulative H_2_O_2_ was also normalized to cell number as determined by the dsDNA microplate assay.

### Quantification of superoxide production with dihydroxyethidium (DHE)

Superoxide production in co- and mono-cultured Hepa1-6 and mHSC, after co-culture model procedures as described above, were quantified by superoxide probe DHE (Fink et al., 2004) and detected by a CLARIOstar fluorometer at λ_Ex_/λ_Em_ = 520-20/590-40 nm in well-scan mode. The DHE assay was performed in a modified Hank’s buffered salt solution for ambient air (130 mM NaCl, 4.56 mM KCl, 1.27 mM CaCl_2_, 0.41 mM MgSO_4_, 0.39 mM MgCl_2_, 0.45 mM NaH_2_PO_4_, 0.34 mM Na_2_HPO_4_, 10 mM HEPES-NaOH [pH 7.4]) at 37°C in ambient air (Chang et al., 2016). Briefly, after removing the spent media, the inserts and wells were washed twice with the experimental medium to remove serum. Inserts and wells were separated to measure superoxide production in a cell type-specific way. At the start of the assay, 120 µL and 520 µL experimental medium was added to all inserts and wells, respectively. The background of all wells and inserts were first scanned and then 5 µM DHE (final concentration; Abcam, #ab145360) was added to each insert and well. After 3 hours of incubation at 37°C, all wells and inserts were scanned again. The background-corrected DHE fluorescence was normalized to cell number as determined by the dsDNA microplate assay.

### Double-stranded DNA (dsDNA) microplate assay

The dsDNA microplate assay was performed based on the work of McCaffery et al. (McCaffrey et al., 1988) and Kapuściński and Skoczylas (Kapuściński & Skoczylas, 1977). Briefly, at the end of experiments, cells in the well or the Transwell insert were fixed with 4% (w/v) PFA for 15 minutes and washed twice with staining buffer (5 mM Na_2_SO_4_ and 5 mM EDTA-NaOH [pH 8.0], sterile-filtered through a 0.2 µm filter unit). For a stoichiometric binding of Hoechst33342 to dsDNA, cells in the wells and inserts were stained overnight with 400 µL and 200 µL 5 µg/mL Hoechst33342 (Sigma, #B2261), respectively. The microplate was scanned in a CLARIOstar fluorometer at λ_Ex_/λ_Em_ = 350-10/490-40 nm. To quantify dsDNA content, a dsDNA standard of 20 µg/mL to 0.3125 µg/mL was constructed from calf thymic dsDNA (Sigma, #D1501) in staining buffer with 5 µg/mL Hoechst33342. To convert dsDNA content to cell numbers, Hepa1-6 and mHSC were seeded at 200,000 to 1563 cells per well in serial 2-fold dilution in 96-well plates, fixed in 4% PFA after attachment, from which the dsDNA content was quantified to calculate the average dsDNA per cell. The dsDNA contents are 7.8 pg/cell and 11.7 pg/cell for Hepa1-6 cells and mHSCs, respectively.

### Statistical analysis

Data analyses and statistical tests were performed using the GraphPad Prism 9 & 10 software and the R 4.4.2 and RStudio software. In all figure legends it is indicated which statistical tests were performed, as well as the p-value and the amount of technical/biological replicates.

## Results

### Hepatic stellate cells increase the proliferation of Hepa1-6 cells and reprogram their metabolism

To investigate the metabolic crosstalk between HSCs and HCC cells, we established a Transwell co-culture model of activated mHSCs and Hepa1-6 cells (Fig. 1A). This setup allowed for rapid isolation of each cell-type after co-culture to study their individual metabolism. Analysis of cell proliferation of Hepa1-6 cells after 72 hours of co-culture with mHSCs showed a 1.8-fold increase in proliferation of Hepa1-6 as compared to mono-cultured cells (Fig. 1B). Importantly, no change in proliferation was observed in mHSCs cultured in the absence or presence of Hepa1-6; and co-culturing Hepa1-6 with Hepa1-6, namely having Hepa1-6 cells in both the well and insert, did not increase Hepa1-6 proliferation in the well (Fig. S1A). This indicates that the increased proliferation of Hepa1-6 in co-culture with mHSC was driven specifically by mHSCs, not merely due to the presence of more cells.

**Figure 1.**
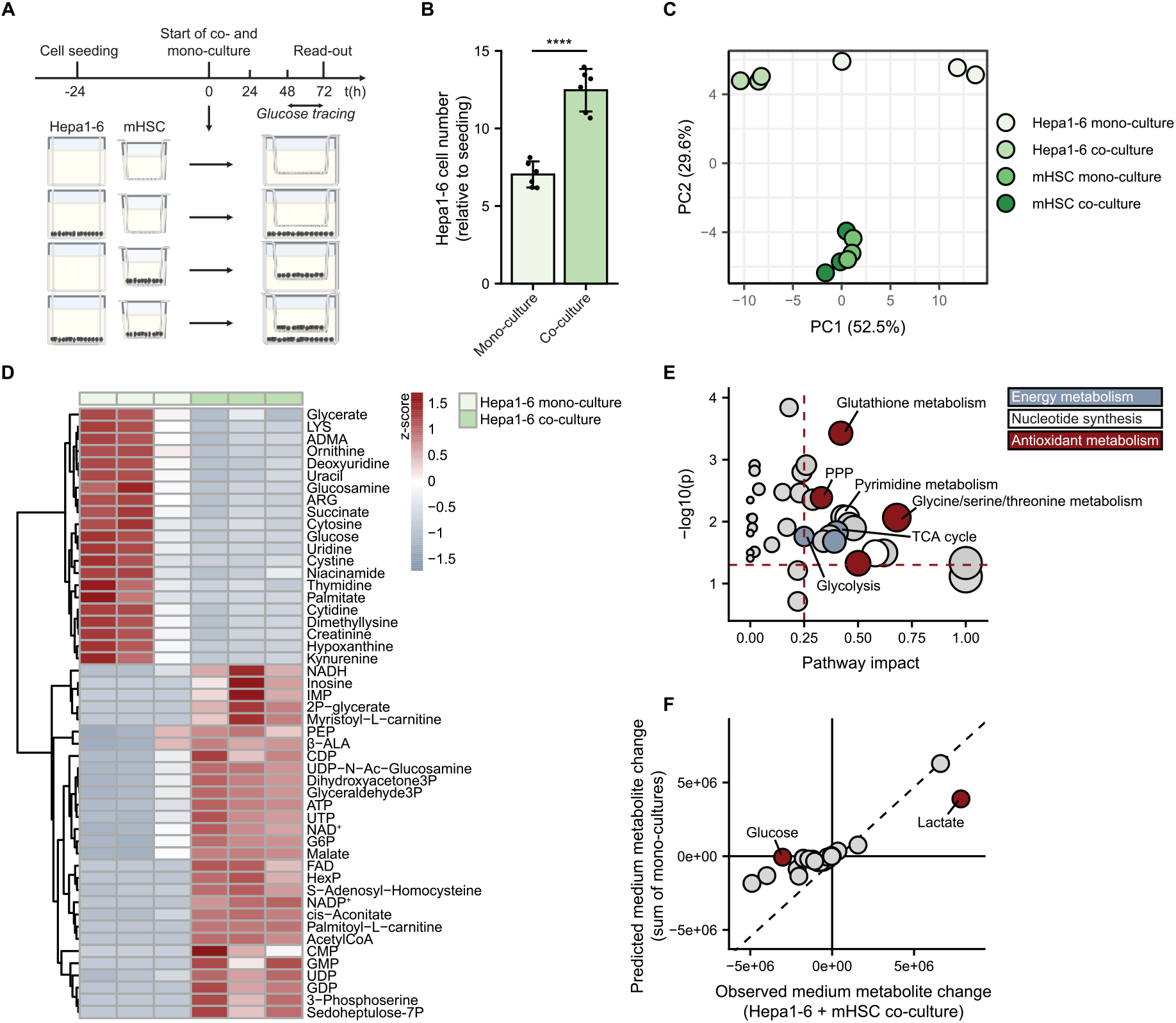
mHSCs increase the proliferation of Hepa1-6 cells and reprogram their metabolism. **A)** Timeline and setup of the Transwell co-culture model. Hepa1-6 cells and mHSCs were seeded 24 hours prior to starting the co- and mono-cultures. Subsequently, cells were cultured for an additional 72 hours, at which point read-outs were performed including proliferation assays and metabolomics analyses. During the final 24 hours of culture, cells were incubated with [U-^13^C_6_]-D-glucose for isotope tracing. **B)** Proliferation rates of co- and mono-cultured Hepa1-6 cells, depicted as fold change of cell number relative to the seeding density. Combined data from two representative independent experiments (n = 3 technical replicates/experiment) are shown. Data are presented as mean ± SD. An unpaired t-test with Welch’s correction was used for statistical analysis, significance is indicated by asterisks (**** = p < 0.0001). **C)** PCA based on total levels of intracellular metabolites of co- and mono-cultured mHSCs and Hepa1-6 cells (n = 3 replicates per condition). **D)** Hierarchical clustering of top 50 differential intracellular metabolites across co- and mono-cultured Hepa1-6 cells. Values are presented as z-scores of total metabolite levels and n = 3 replicates per condition were included. **E)** Pathway impact analysis of all identified intracellular metabolites, comparing co- and mono-cultured Hepa1-6 cells using MetaboAnalyst (n = 3 replicates per condition). Pathway impact is depicted on the x-axis and significance is illustrated on the y-axis. Highlighted are pathways with p < 0.05 and a pathway impact ≥ 0.25 related to energy metabolism (blue), nucleotide synthesis (white) and antioxidant metabolism (red). **F)** Uptake or secretion of identified extracellular metabolites in co-culture (observed medium metabolite change) versus the summed uptake or secretion in mHSC and Hepa1-6 mono-cultures (predicted medium metabolite change) (n = 3 replicates per condition). Metabolites are visualized as dots, with those meeting the following two criteria annotated and highlighted in red: a significant difference between observed and predicted values (p < 0.05, calculated with an unpaired t-test) and an absolute horizontal distance to the diagonal line greater than 1.5e+06, based on the absolute difference between observed and predicted values.

To investigate whether the increased proliferation of co-cultured Hepa1-6 cells was accompanied with specific metabolic changes, we performed LC-MS-based semi-targeted metabolomics on both intracellular and extracellular metabolites after 72 hours of (co-)culture, with [U-^13^C_6_]-D-glucose added during the final 24 hours (Fig. 1A). A principle component analysis (PCA) of total intracellular metabolite levels revealed a clear distinction between Hepa1-6 and mHSC, highlighting their intrinsically distinct metabolic signatures (Fig. 1C). While co- and mono-cultured mHSCs clustered together, co- and mono-cultured Hepa1-6 cells separated along PC1 (52.5% variability) (Fig. 1C), indicating that co-culture with mHSCs impacts the metabolome of Hepa1-6 cells but not vice versa. In line with this, hierarchical clustering analysis of all intracellular metabolites and the top 50 differential metabolites showed specific clusters of up- and downregulated metabolites across co- and mono-cultured Hepa1-6 cells (Fig. S1B, Fig. 1D). Differences were less pronounced in mono- and co-cultured mHSCs (Fig. S1B). A pathway impact analysis comparing co- versus mono-cultured Hepa1-6 showed that co-culture significantly altered energy metabolism (e.g., glycolysis, the tricarboxylic acid (TCA) cycle), nucleotide synthesis (e.g., pyrimidine metabolism) and metabolic pathways involved in modulating oxidative stress (e.g., glutathione metabolism, serine metabolism, and the pentose phosphate pathway (PPP)) (Fig. 1E). Pathway impact analysis comparing co- and mono-cultured mHSCs yielded only few regulated pathways, of which glutathione metabolism was most significantly impacted (Fig. S1C). To investigate whether co-culture affects metabolite uptake or secretion via cell-cell interactions, we compared the medium metabolite changes of Hepa1-6-mHSC co-culture to the sum of those of Hepa1-6 and mHSC mono-cultures (Fig. 1F). In the absence of HCC-HSC interaction effects, metabolite changes are expected to be the same and the data point would fall on the diagonal line. Based on statistical significance (p < 0.05) and an absolute horizontal distance to the diagonal line exceeding 1.5 x 10^6^, this analysis revealed cell-cell interactions affected glucose and lactate levels. Co-culture of Hepa1-6 cells and mHSCs synergistically increased the uptake of glucose and secretion of lactate (Fig. 1F), as also illustrated by their absolute levels in the medium (S1D), suggesting an increase in glycolysis. In line with this, the glucose-tracing experiment showed that co-cultured Hepa1-6 cells increased incorporation of glucose-derived carbon in multiple metabolites, including those in glycolysis, the TCA cycle, PPP, serine synthesis pathway (SSP), nucleotide synthesis and glutathione metabolism (Fig. S1E). Together, these results demonstrate that in Hepa1-6-mHSC co-culture, mHSCs promote Hepa1-6 proliferation and rewired glucose metabolism to support energy production, antioxidant, and biosynthetic pathways in Hepa1-6 cells.

### mHSC-induced proliferation of Hepa1-6 cells is dependent on the uptake of glucose, which is used in both energy and biosynthetic pathways

As mHSCs rewired glucose metabolism in Hepa1-6, which included enhanced glucose uptake and lactate secretion in co-culture, we next investigated whether this metabolic rewiring contributed to the increased proliferation of Hepa1-6 observed in co-culture. To this end, we performed proliferation assays after co- and mono-culturing in the presence of either 2-DG, a direct inhibitor of glycolysis, or BAY-876, an inhibitor of GLUT1. Administration of a low dose of either 2-DG or BAY-876 significantly decreased the proliferation rate of co-cultured Hepa1-6 cells, reducing these to mono-culture levels (Fig. 2A). In contrast, the proliferation rates of mono-cultured Hepa1-6 remained unaffected by the same low-dose 2-DG or BAY-876. These findings together indicate that enhanced glucose uptake and glycolytic metabolism contribute specifically to the increased proliferation of co-cultured Hepa1-6 cells.

**Figure 2.**
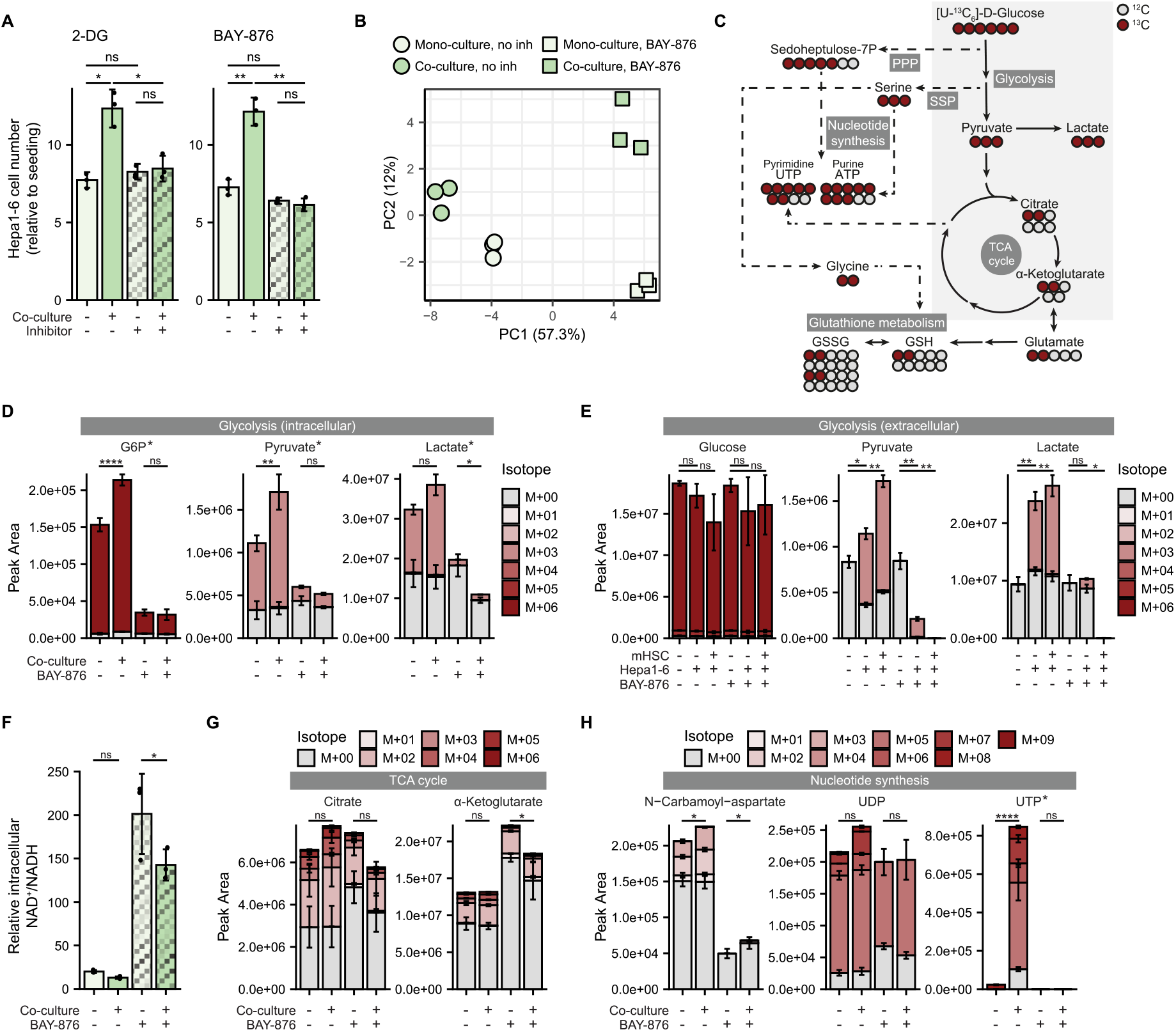
mHSC-induced proliferation of Hepa1-6 cells is dependent on the uptake of glucose, which is used in both energy and biosynthetic pathways. For panels **B** and **D-H**, metabolomics data were obtained from co- and mono-cultured Hepa1-6 cells after a 72-hour incubation with [U-^13^C_6_]-D-glucose in the presence or absence of BAY-876 (5 nM) (n = 3 replicates per condition). **A)** Proliferation rates of co- and mono-cultured Hepa1-6, depicted as fold change of cell number relative to the seeding density, in the absence or presence (72 hours) of a glycolysis inhibitor (2-DG, 0.1 mM, left panel) or a GLUT1 inhibitor (BAY-876, 5 nM, right panel) (n = 3 technical replicates per condition). Data are presented as mean ± SD. An unpaired t-test with Welch’s correction was used for statistical analysis, significance is indicated by asterisks (* = p < 0.05, ** = p < 0.005). **B)** PCA based on total ^13^C-labeled fractions of intracellular metabolites. **C)** Schematic diagram of the potential incorporation of labeled carbons from [U-^13^C_6_]-D-glucose in downstream metabolites and pathways. Metabolites in glycolysis, the TCA cycle, the PPP and SSP, nucleotide synthesis and glutathione metabolism pathways are visualized. **D, G, H)** Levels and isotopologue distribution patterns of intracellular glycolysis metabolites (**D**), TCA cycle metabolites (**G**), and pyrimidines (**H**). Data are presented as mean ± SD. Statistical significance was assessed using a two-way ANOVA, which showed a significant interaction between culture (mono-/co-culture) and inhibitor (no inhibitor/BAY-876) conditions for metabolites labeled with an asterisk, followed by Šídák-adjusted post hoc multiple comparisons based on estimated marginal means. Significance shown in the figure corresponds to these post hoc comparisons, illustrated by asterisks (* = p < 0.05, ** = p < 0.01, **** = p < 0.0001). **E)** Levels and isotopologue distribution patterns of extracellular glycolysis metabolites across experimental conditions, including a medium-only control. Data are presented as mean ± SD. Statistical significance was determined using unpaired t-tests with Welch’s correction and with Benjamini–Hochberg correction for multiple testing. Significance is indicated by asterisks for adjusted p-values (* = adjusted p < 0.05, ** = adjusted p < 0.01). **F)** Relative NAD^+^/NADH ratio. Values were calculated using total peak areas of NAD^+^ and NADH as assessed by metabolomics. Data are presented as mean ± SD. Statistical significance was assessed using a two-way ANOVA, which showed no significant interaction between culture (mono-/co-culture) and inhibitor (no inhibitor/BAY-876) conditions (p = 0.109), followed by Šídák-adjusted post hoc multiple comparisons based on estimated marginal means. Significance shown in the figure corresponds to these post hoc comparisons, illustrated by asterisks (* = p < 0.05).

To investigate which downstream metabolic pathways could be involved in driving the increased proliferation under co-culture conditions, we again performed [U-^13^C_6_]-D-glucose tracing in mono-and co-cultures, now for 72 hours in the absence or presence of BAY-876 (Fig. S2A). Compared to the previous tracing setup, this approach avoided an additional medium change that could interfere with inhibitor efficacy, while potentially capturing alterations in pathway activity arising during initial stages of co-culture. PCA based on the fraction of ^13^C incorporation across metabolites revealed that BAY-876-treated cultures separated from control groups, indicating that BAY-876 induces a distinct metabolic phenotype in both mono- and co-cultured Hepa1-6 cells (Fig. 2B). We next assessed ^13^C incorporation in pathways identified in our initial metabolomics screen in more detail (Fig. 2C). As expected, 72 hours of co-culture increased the total levels and ^13^C-labeled fractions of glycolytic intermediates (G6P, pyruvate, lactate) in Hepa1-6 (Fig. 2D) and increased the secretion of (glucose-derived) pyruvate and lactate (Fig. 2E), all of which is indicative of higher glycolytic activity. BAY-876 treatment induced a severe downregulation in total levels of intracellular and extracellular glycolysis intermediates and almost completely abolished label incorporation in these metabolites in both mono- and co-cultured cells (Fig. 2D, Fig. 2E). Moreover, BAY-876 treatment decreased overall levels of NADH, abolishing differences between co- and mono-cultured Hepa1-6 cells and greatly increased the NAD^+^/NADH ratio in both cultures (Fig. S2B, 2F). This likely reflects the lack of glycolytic activity in BAY-876-treated cells due to inhibition of glucose uptake. Thus, BAY-876 treatment diminished differences in glycolysis between co- and mono-cultured Hepa1-6 cells, confirming that glycolysis is a primary target of BAY-876 and likely necessary for the increased proliferation of co-cultured Hepa1-6 cells.

Further downstream, the levels of and ^13^C incorporation in the PPP (e.g., sedoheptulose-7P), SSP (e.g., serine) and TCA cycle metabolites (e.g., citrate, α-ketoglutarate) were relatively unaffected by co-culture in the absence of BAY-876 (Fig. S2C, 2G), although the fractions of labelled serine increased (Fig. S2C). Both glutathione (GSH) and oxidized glutathione (GSSG) showed increased labeling from [U-^13^C_6_]-D-glucose in co-cultured compared to mono-cultured Hepa1-6 cells, suggestive of increased glutathione synthesis from glucose presumably via the SSP (Fig. S2D). BAY-876 treatment inhibited the PPP as evidenced by the dramatic reduction in sedoheptulose-7P levels. In contrast, it had a modest effect on the SSP, TCA cycle and glutathione synthesis (Fig. S2C, 2G, S2D). While BAY-876 treatment lowered [U-^13^C_6_]-D-glucose labeling in TCA cycle metabolites and (oxidized) glutathione in mono- and co-cultures, unlabeled levels of these metabolites remained stable or even increased (Fig. 2G, S2D). Likely, Hepa1-6 cells switch to other carbon sources (e.g., glutamine) to compensate for the lack of glucose. Moreover, this suggests that BAY-876 does not exert its growth inhibitory effects through these pathways. Notably, in the absence of BAY-876, co-culture enhanced the ^13^C labeling of pyrimidine precursors (e.g., N-carbamoyl-aspartate) and also increased the fractions of pyrimidine nucleotides [M>5] (e.g., UDP, UTP), both of which are indicative of enhanced *de novo* pyrimidine synthesis in co-cultured Hepa1-6 cells (Fig. 2H). Purine nucleotides did not show increased levels and relative ^13^C incorporation in co- versus mono-cultured Hepa1-6 (Fig. S2E). BAY-876 treatment decreased the pyrimidine precursor and nucleotide labeling, including [M>5] fractions, and brought levels in co-cultured cells back to mono-culture levels (Fig. 2H), suggesting BAY-876 inhibits *de novo* pyrimidine synthesis. Notably, BAY-876 also diminished ATP levels in co-cultured cells specifically (Fig. S2E), suggesting these cells suffer from a lack of energy.

Together, these data confirm that co-culture with mHSCs induces profound changes in glucose uptake and metabolism and increases glucose usage in pathways that drive tumor growth and proliferation, including glycolysis, glutathione synthesis and *de novo* pyrimidine synthesis in Hepa1-6 cells. Targeting glucose metabolism in co-cultured Hepa1-6 cells severely reduced glycolysis and PPP intermediates, and brought pyrimidine nucleotide levels in co-cultured cells back to mono-culture levels. Our results suggest that mHSCs modulate glycolysis, PPP activity and/or *de novo* pyrimidine synthesis in Hepa1-6 cells to sustain the increased proliferation of Hepa1-6 cells in co-culture.

### mHSCs induce HIF signaling in Hepa1-6 cells

To investigate the underlying mechanisms that drives the enhanced proliferation and altered glucose metabolism in co-cultured Hepa1-6 cells, we performed RNA sequencing on co- and mono-cultures. PCA revealed a separation between co- and mono-cultured Hepa1-6 along PC1 (23% variability) (Fig. 3A), indicating that co-culture with mHSCs affected gene expression in Hepa1-6. Differential expression analysis revealed significantly up- and downregulated genes in co-cultured Hepa1-6, with the upregulated genes outnumbering the downregulated genes by ∼2.5-fold when applying an absolute log2FC threshold of 0.25 (Fig. 3B). Pathway enrichment analysis of these significantly upregulated genes in co-cultured Hepa1-6 demonstrated that 7 of the top 8 most enriched terms were related to metabolic pathways (Fig. 3C). In line with our metabolomics findings, these included glycolysis, central carbon metabolism, the PPP and other glucose-related pathways. Additionally, the HIF-1 signaling pathway was identified as the most significantly enriched pathway. Hierarchical clustering of genes involved in HIF signaling indeed revealed upregulation of *Hif-1α* in co-cultured Hepa1-6 cells together with a specific cluster of HIF-1 target genes, including *Vegfa*, *Slc2a1* (encoding GLUT1) and all glycolytic enzymes (Fig. 3D). No upregulation of HIF-related genes was observed in co-cultured mHSCs (Fig. S3A). To confirm the upregulation of HIF signaling in Hepa1-6 cells by mHSCs, we next investigated the nuclear translocation of HIF in mono- and co-cultured Hepa1-6 cells by IF staining and confocal microscopy. When co-cultured with mHSCs, Hepa1-6 cells demonstrated significantly higher nuclear HIF-1α levels than mono-cultured Hepa1-6 cells, with levels increasing to almost those observed in cells treated with DMOG, a known stabilizer of HIF (Fig. 3E). Control conditions, in which we ‘co-cultured’ Hepa1-6 cells with themselves in both the well and the Transwell insert, did not show any increase in HIF-1α nuclear localization. The difference in nuclear HIF-1α levels between Hepa1-6 cells co-cultured with mHSCs and those ‘co-cultured’ with Hepa1-6 demonstrated that mHSCs did not increase nuclear HIF-1α in Hepa1-6 by competing for oxygen in co-culture, but by a mHSC-specific mechanism (Fig. 3E). Taken together, these data show that co-culture with mHSCs increased HIF signaling in Hepa1-6 cells, which is accompanied by increased cell proliferation and a rewiring of glucose metabolism.

**Figure 3.**
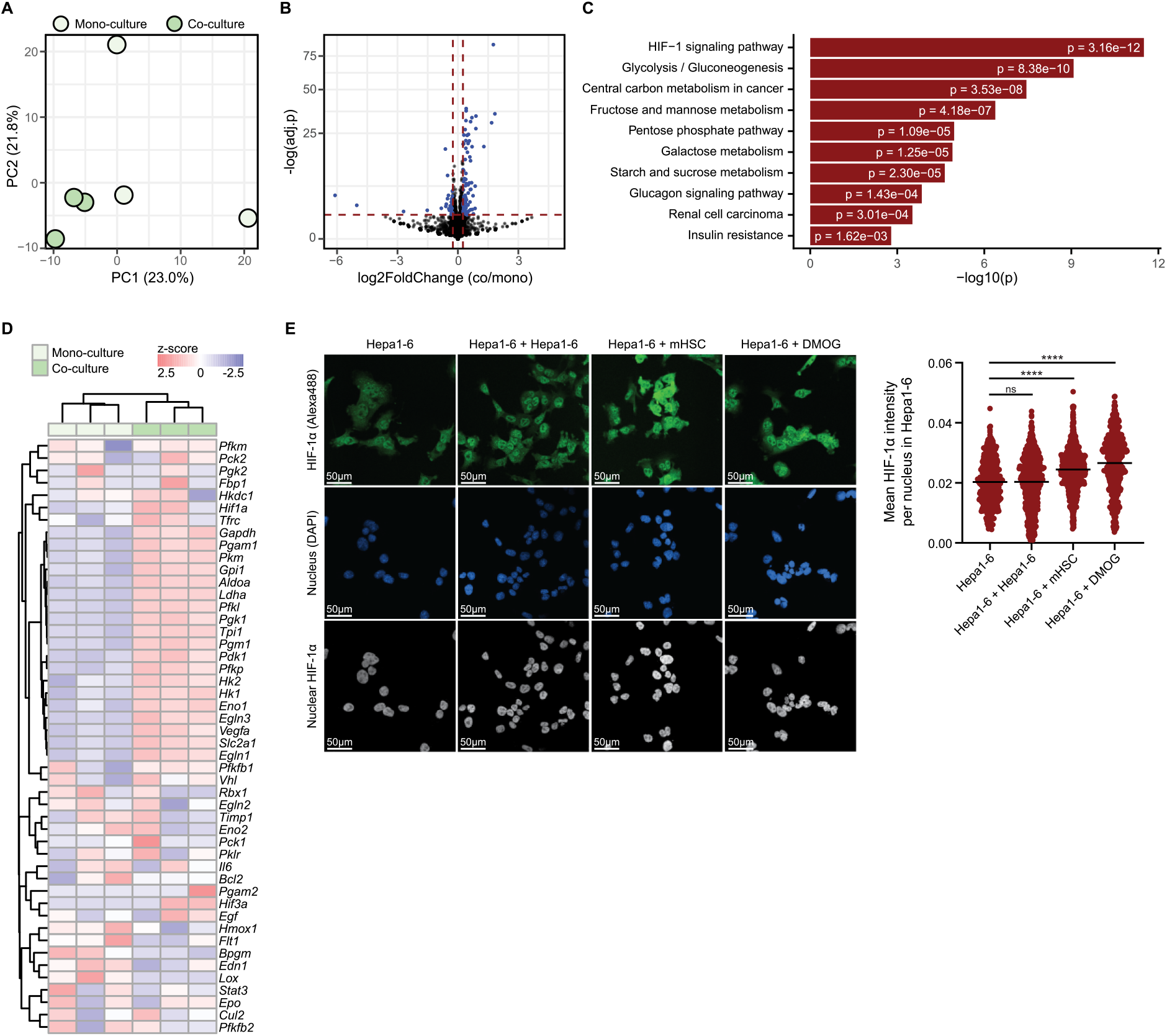
mHSCs induce HIF signaling in Hepa1-6 cells. RNA sequencing data of Hepa1-6 cells, obtained after 72 hours of mono- or co-culture with mHSC cells. Raw readcounts were normalized using The Utrecht University ATWiCS ShinyApp. **A)** PCA based on normalized gene expression levels of co- and mono-cultured Hepa1-6 cells (n = 3 replicates per condition). **B)** Volcano plot showing the differentially expressed genes in co-cultured versus mono-cultured Hepa1-6 cells. Log2-transformed fold change is depicted on the x-axis, adjusted p-value is indicated on the y-axis. Identified genes are indicated with black dots, blue dots indicate genes that are significantly different expressed (absolute log2FC ≥ 0.25 and adjusted p-value < 0.05) between co- and mono-cultured Hepa1-6 cells. **C)** Gene set enrichment and pathway analysis of significantly upregulated genes in co-cultured Hepa1-6 (illustrated in 3B, log2FC ≥ 0.25, adjusted p-value < 0.05), using Enrichr. The bar chart is showing the top 10 enriched pathways from the KEGG2021 gene set library, based on the −log10(p-value) with the actual p-value are shown next to each term. The pathway at the top with the smallest p-value has most significant overlap with the input gene set. All pathways presented are also significant based on adjusted p-value (< 0.05). **D)** Hierarchically clustered heatmap showing z-score-normalized gene expressions of genes related to HIF signaling in co- and mono-cultured Hepa1-6 based (n = 3 replicates per condition). **E)** Left: representative microscopy confocal images of the immunofluorescence staining of HIF-1α (Alexa488) (top panels) and the nucleus (DAPI) (middle panels) depicted per condition, which include Hepa1-6 mono-cultured, Hepa1-6 ‘co-cultured’ with Hepa1-6, Hepa1-6 co-cultured with mHSCs and Hepa1-6 ‘mono-cultured’ with DMOG. Bottom panels show nuclear HIF-1α signal obtained using a DAPI-derived nuclear mask. Right: the nuclear HIF-1α intensity is quantified and depicted as mean intensity per nucleus and per condition. Each dot represents a cell that is quantified for its signal, with the horizontal bar representing the mean. Five separate (2 x 2 large) images were included for analysis per condition. Statistical testing included Games-Howell’s multiple comparisons test, with significance indicated by asterisks (**** = adjusted p < 0.0001).

Inhibition of HIF signaling reverses mHSC-induced metabolic reprogramming and proliferation in Hepa1-6 cells To examine if the increased HIF signaling in co-culture was necessary for the enhanced proliferation of Hepa1-6 cells, we suppressed HIF-1α and HIF-2α signaling with PX-478 and PT-2385, respectively (Koh et al., 2008; E. M. Wallace et al., 2016), in co- and mono-cultured Hepa1-6. Administration of either of these two agents induced a significant decrease in proliferation of co-cultured Hepa1-6, abolishing the difference in growth between co- and mono-cultures (Fig. 4A). To assess whether HIF inhibition also reversed the metabolic rewiring observed after co-culture, we performed [U-^13^C_6_]-D-glucose tracing in the presence of PT-2385. As expected, PCA based on total ^13^C-labeled fractions of intracellular metabolites revealed a clear separation between untreated and PT-2385-treated cells along PC2 (21.8% variability) (Fig. 4B). Like BAY-876, PT-2385 downregulated both the levels of and ^13^C incorporation in glycolysis metabolites in both mono- and co-cultured cells (Fig. 4C), largely eliminating differences between co- and mono-culture. This suggests that HIF inhibition counteracts the increase in glycolysis induced by co-culture, although to a lesser extent than BAY-876. As compared to GLUT1 inhibition by BAY-876, inhibiting HIF-2α transcriptional activity with PT-2385 showed less pronounced effects on the TCA cycle, PPP, SSP and glutathione synthesis (Fig. 4D, S4A, S4B), indicating that residual transporter and enzyme activities are sufficient to maintain glucose entry into these pathways. The *de novo* pyrimidine nucleotide synthesis was partly inhibited by PT-2385, as evidenced by reduced levels and labeling in N-carbamoyl-aspartate (Fig. 4E). However, the levels of most nucleotides remained stable or increased (Fig. 4E, S4C), suggesting that nucleotide pools were maintained despite HIF-2α inhibition. Isotopologue distributions also revealed limited changes, which suggests that *de novo* nucleotide synthesis is not affected by HIF inhibition to a similar extent as by BAY-876 treatment. In line with the decrease in glycolytic activity, as also observed upon BAY-876 treatment, PT-2385 decreased overall levels of NADH and increased the NAD^+^/NADH ratio in both cultures (Fig. S4D, 4F). Collectively, these findings illustrate that HIF signaling is essential for the increased proliferation rates observed in co-cultured Hepa1-6 and suggest that this effect is at least in part mediated by the inhibition of glycolysis.

**Figure 4.**
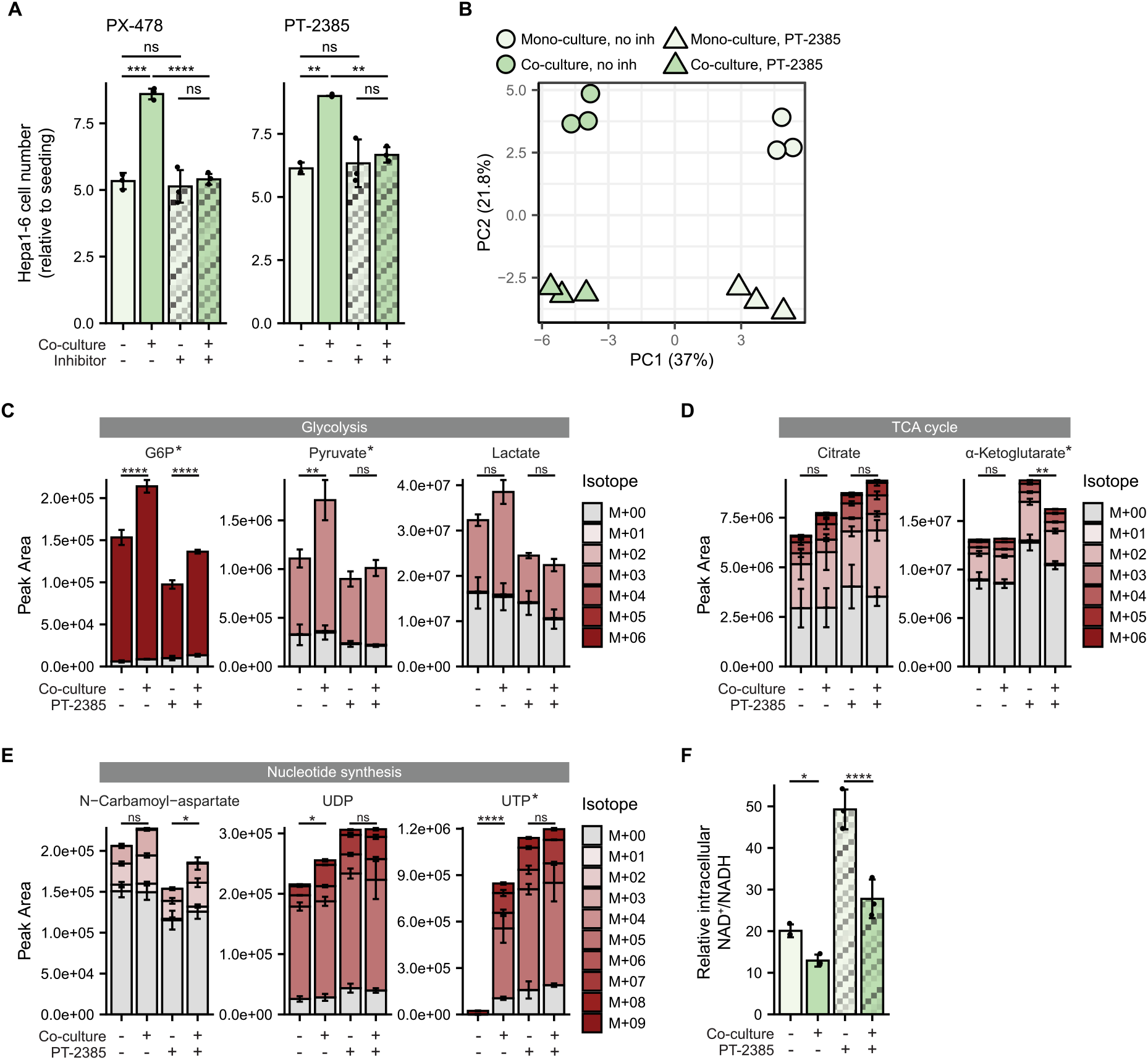
Inhibition of HIF signaling reverses mHSC-induced metabolic reprogramming and proliferation in Hepa1-6 cells. For panels **B-F**, metabolomics data were obtained from co- and mono-cultured Hepa1-6 cells after a 72-hour incubation with [U-^13^C_6_]-D-glucose in the presence or absence of PT-2385 (10 µM) (n = 3 replicates per condition). NB: PT-2385 and BAY-876 conditions were combined in one metabolomics experiment. Control (untreated) co- and mono-cultured Hepa1-6 cells are therefore identical to those presented in Fig. 2 and S2. **A)** Proliferation rates of co- and mono-cultured Hepa1-6 cells, depicted as fold change of cell number relative to the seeding density, in the absence or presence (72 hours) of a suppressing agent of HIF-1α (PX-478, 5 µM, left panel) or a HIF-2α inhibitor (PT-2385, 10 µM, right panel) (n = 3 technical replicates per condition). Data are presented as mean ± SD. An unpaired t-test with Welch’s correction was used for statistical analysis, significance is indicated by asterisks (** = p < 0.01, *** = p < 0.0005, **** = p < 0.0001). **B)** PCA based on total ^13^C-labeled fractions of intracellular metabolites. **C, D, E)** Levels and isotopologue distribution patterns of intracellular glycolysis metabolites (**C**), TCA cycle metabolites (**D**) and pyrimidines (**E**). Data are presented as mean ± SD. Statistical significance was assessed using a two-way ANOVA, which showed a significant interaction between culture (mono-/co-culture) and inhibitor (no inhibitor/ PT-2385) conditions for metabolites labeled with an asterisk, followed by Šídák-adjusted post hoc multiple comparisons based on estimated marginal means. Significance shown in the figure corresponds to these post hoc comparisons, illustrated by asterisks (* = p < 0.05, ** = p < 0.01, **** = p < 0.0001). **F)** Relative NAD^+^/NADH ratio. Values were calculated using total peak areas of NAD^+^ and NADH as assessed by metabolomics. Data are presented as mean ± SD. Statistical significance was assessed using a two-way ANOVA, which showed a significant interaction between culture (mono-/co-culture) and inhibitor (no inhibitor/PT-2385) conditions (p = 0.007), followed by Šídák-adjusted post hoc multiple comparisons based on estimated marginal means. Significance shown in the figure corresponds to these post hoc comparisons, illustrated by asterisks (* = p < 0.05, **** = p < 0.0001).

### mHSC-induced ROS is a potential crosstalk mediator that can promote HIF activation in Hepa1-6 cells

Finally, we questioned how Hepa1-6-mHSC crosstalk induced HIF signaling in Hepa1-6 cells. ROS is a well-known inducer of HIF, and in the datasets described above we found several indications that ROS could be involved in activating HIF signaling in our Hepa1-6-mHSC co-culture system. First, glutathione metabolism was identified as the top regulated pathway in co-cultured Hepa1-6 cells and mHSCs (Fig. 1E, S1C). Second, a trend towards an increased GSSG/GSH ratio was observed in co-cultured Hepa1-6 cells compared to mono-cultured cells, obtained from the 24-hour [U-^13^C_6_]-D-glucose tracing data, indicative of intracellular oxidative stress (Fig. S5A). Based on the 72-hour [U-^13^C_6_]-D-glucose tracing data, this tendency was reversed by PT-2385 but not BAY-876 treatment, suggesting an association with HIF signaling (Fig. S5B). Third, oxidative stress-related genes were differentially expressed in mHSCs upon co-culture. Expression of the *catalase* gene (*Cat*) and *superoxide dismutase* genes (*Sod1/2/3*), encoding enzymes involved in scavenging hydrogen peroxide (H_2_O_2_) and superoxide, respectively, were downregulated in co-cultured mHSC cells as compared to monoculture (Fig 5A). Superoxide is a highly reactive free radical and thus has a very short half-life and only reacts locally, which is unlikely to be a messenger on its own. However, superoxide dismutase can convert superoxide into H_2_O_2_, which is a non-radical ROS with much longer half-life and can therefore travel a longer distance, diffuse across cell membranes and function as a crosstalk molecule. Based on the above-mentioned data and reasoning, we hypothesized that a downregulation of catalase in mHSCs could lead to increased extracellular levels of H_2_O_2_, in turn causing oxidative stress and activation of HIF signaling in Hepa1-6 cells (Fig. 5B).

**Figure 5.**
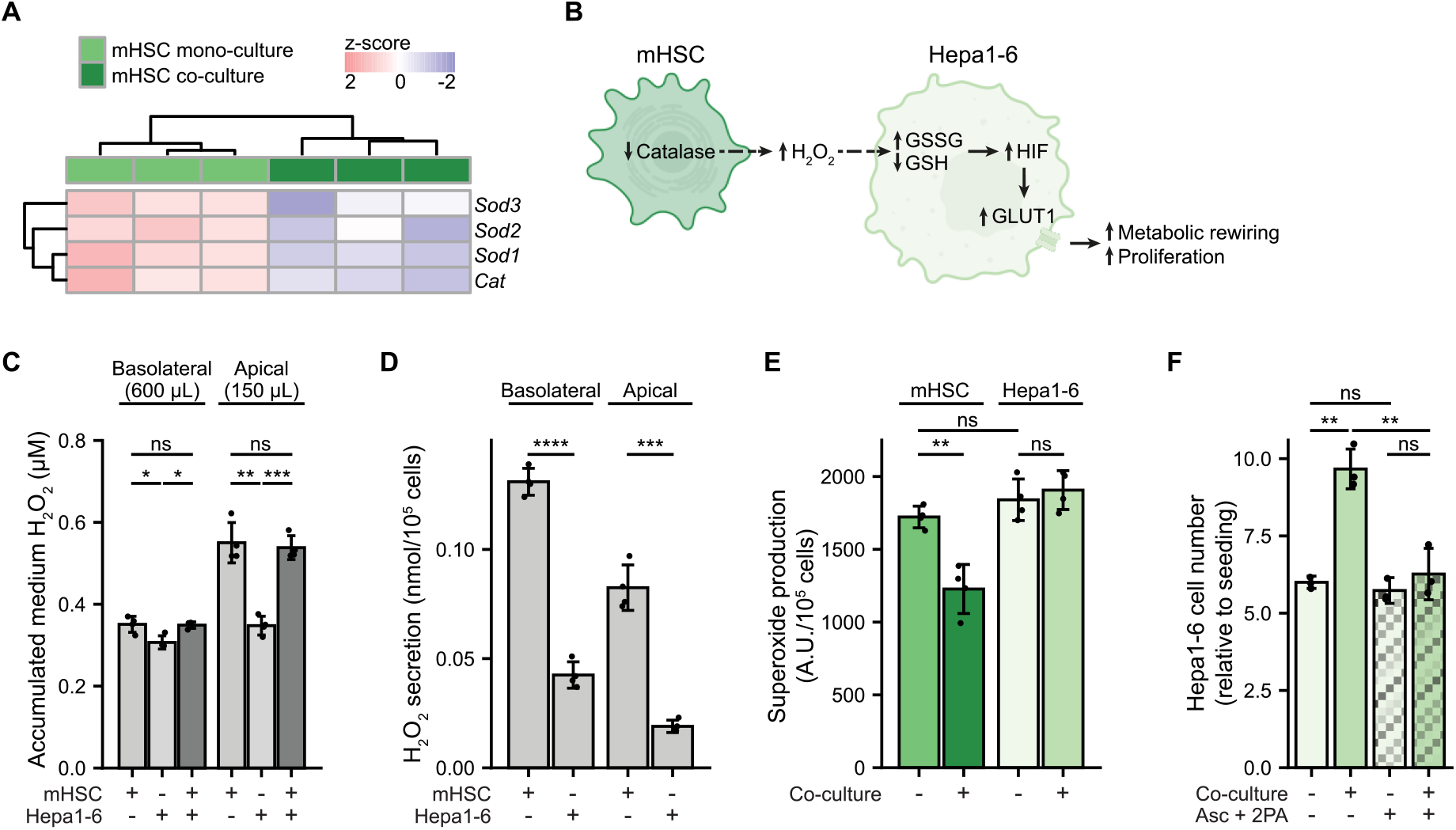
mHSC-induced ROS is a potential crosstalk mediator that can promote HIF activation in Hepa1-6 cells. **A)** Hierarchically clustered heatmap showing z-score-normalized expression of genes encoding catalase (*Cat*) and superoxide dismutase 1/2/3 (*Sod1/2/3*) in co- and mono-cultured mHSCs (n = 3 per condition) (these data are obtained from the same transcriptomics experiment as presented in Fig. 3 and S3). **B)** Working model of our findings, illustrating mHSC (dark green) and Hepa1-6 (light green) and the potential role of ROS in the activation of HIF in co-culture. **C)** Accumulated H_2_O_2_ secretion (µM) over 18 hours in medium of the basolateral compartment (bottom well) with 600 µL volume and the apical compartment (insert) with 150 µL volume, depicted for mHSC and Hepa1-6 mono-cultures and co-culture, with each n = 4 replicates. Data are presented as mean ± SD. Statistical tests included Dunnett’s T3 multiple comparisons, which is visualized in the graph (* = adjusted p < 0.05, ** = adjusted p < 0.005, *** = adjusted p < 0.0005), after a Brown-Forsythe and Welch ANOVA test which also indicated significance across groups. **D)** H_2_O_2_ secretion in nmol per 10^5^ cells over 18 hours, calculated based on mHSC and Hepa1-6 mono-cultures from panel C with each n = 4 replicates, depicted for both the basolateral compartment (bottom well) and the apical compartment (insert). Data are presented as mean ± SD. Statistical tests to calculate significance were done using unpaired t-test with Welch’s correction (*** = p < 0.001, **** = p < 0.0001). **E)** ROS production rate in arbitrary units per 10^5^ cells, depicted for mHSC mono- and co-cultured with Hepa1-6, and for Hepa1-6 mono- and co-cultured with mHSC, with each n = 4 replicates. Data are presented as mean ± SD. Statistical significance between conditions was calculated using an unpaired t-test (** = p < 0.005). **F)** Proliferation rates of co- and mono-cultured Hepa1-6 cells, depicted as fold change of cell number relative to the seeding density, in the absence or presence (72 hours) of a stabilized vitamin C solution containing 75 µM ascorbate (Asc) and 500 µM 2-phosphate-L-ascorbate (2PA) (n = 3 technical replicates per condition). Data are presented as mean ± SD. An unpaired t-test with Welch’s correction was used for statistical analysis, significance is indicated by asterisks (** = p < 0.01).

To test this hypothesis, we measured H_2_O_2_ secretion by mHSCs and Hepa1-6 cells in mono- and co-cultures. Mono-cultured mHSCs and Hepa1-6 cells both secreted H_2_O_2_, resulting in H_2_O_2_ accumulation in both the basolateral compartment (600 µL) containing Hepa1-6 cells and the apical compartment (150 µL) containing mHSCs (Fig. 5C). In the co-culture of Hepa1-6 cells and mHSCs, H_2_O_2_ accumulation in both apical and basolateral compartments was higher than Hepa1-6 mono-culture levels and comparable to mHSC mono-culture levels. As a result, the accumulated H_2_O_2_ secretion in co-culture medium was about half of the expected sum of Hepa1-6 and mHSC mono-cultures, indicating that a portion of medium H_2_O_2_ was neutralized by cells in co-culture (Fig. 5C). Using these H_2_O_2_ accumulation values, we next analyzed the H_2_O_2_ secretion rate per cell based on compartment volumes. mHSCs emerged as the major source of extracellular H_2_O_2_, as they secreted significantly more H_2_O_2_ per cell compared to Hepa1-6 (Fig. 5D). We next asked if intracellular superoxide production was the source of H_2_O_2_ secretion. To this end, we used dihydroethidium to quantify superoxides (Fink et al., 2004). Mono-cultured Hepa1-6 and mHSC showed similar superoxide production per cell (Fig. 5E), despite the higher contribution of mHSCs to extracellular H_2_O_2_ levels compared to Hepa1-6. In fact, co-cultured mHSCs demonstrated lower superoxide production compared to their mono-cultured counterparts. This suggests that an alternative source of H_2_O_2_ or a decrease in H_2_O_2_ scavenging drive H_2_O_2_ secretion by mHSCs under co-culture conditions, in line with our transcriptomic findings showing decreased expression of *Cat* and *Sod1-3* (Fig. 5A). Superoxide production was not affected in co-cultured Hepa1-6 compared to mono-cultured cells (Fig. 5E). These findings together suggest that Hepa1-6 cells do not substantially contribute to increased ROS production under co-culture conditions but instead rely on mHSCs as a source of H_2_O_2_, through a mechanism other than superoxide production.

Finally, to assess the importance of oxidative stress for the HIF-enhanced proliferation of co-cultured Hepa1-6, we supplemented mono- and co-cultures with a mixture of ascorbate (Asc) and 2-phosphate-L-ascorbate (2PA) to maintain intracellular vitamin C levels over the 72-hour period of co-culture (Frikke-Schmidt & Lykkesfeldt, 2010). Vitamin C is an antioxidant known to reduce ROS and subsequent oxidative stress, and to inhibit HIF signaling by promoting HIF-1/2α degradation. Supplementation of this Asc + 2PA mixture significantly decreased the proliferation of co-cultured Hepa1-6 cells, abolishing the growth difference between co- and mono-cultures (Fig. 5F), while mono-cultures were unaffected.

Taken together, our data demonstrate that mHSCs rewire Hepa1-6 metabolism through activation of HIF and upregulation of glucose metabolism. This glycolytic switch drives an increase in proliferation of Hepa1-6 cells. We show that oxidative stress plays a role in the proliferative phenotype of co-cultured Hepa1-6 cells and identify mHSC-derived H_2_O_2_ as a potential crosstalk mediator that can promote ROS-dependent HIF activation. By elucidating this HSC-dependent crosstalk as a potential mechanism of HIF activation in HCC, a key driver of angiogenesis and target of current therapies, our findings provide a novel angle for targeting tumor angiogenesis in HCC.

## Discussion

An altered metabolism is a hallmark of cancer, with metabolic crosstalk in the TME representing promising targets that can be exploited in precision-based HCC therapies (Pavlova et al., 2022; Yang et al., 2024). In this study, we model the fibrotic HCC TME by co-culturing mHSCs and Hepa1-6 cells in the Transwell system, and demonstrate that mHSCs interact with Hepa1-6 cells and promote the proliferation of Hepa1-6 cells by enhancing glucose uptake and metabolic rewiring. We further identify HIF signaling as the underlying mechanism that drives both the metabolic rewiring and increased proliferation in Hepa1-6 cells and show that both HIF signaling and glucose uptake can be targeted to counteract this phenotype. Our findings suggest that ROS may contribute to the enhanced proliferation of Hepa1-6 cells under co-culture conditions and identify mHSC-derived H_2_O_2_ as a candidate crosstalk signal that can activate HIF signaling in Hepa1-6. While previous research has identified oxidative stress, HIF signaling and metabolic rewiring as important processes in HCC progression, it has not been fully elucidated how HSC-HCC crosstalk influences these processes and their potential interplay (H. Ma et al., 2018; Sun et al., 2021). Our work identifies HSC-HCC metabolic crosstalk as a novel mechanism through which the TME modulates HCC progression.

We found that when co-cultured with mHSCs, Hepa1-6 cells rewired their metabolism and showed increased activity in glycolysis, the SSP, glutathione metabolism and nucleotide synthesis. This was evident from increased glucose uptake and lactate secretion, and increased glucose-derived carbon incorporation in metabolites part of these pathways, often accompanied by increased total levels. Notably, increased activity in the PPP and TCA cycle was evident from the 24-hour [U-^13^C_6_]-D-glucose tracing, whereas the 72-hour [U-^13^C_6_]-D-glucose tracing did not reveal substantial changes, potentially due to a saturation of glucose utilization with this longer tracing period. Inhibition of glycolysis with 2-DG or GLUT1 by BAY-876 decreased proliferation of co-cultured Hepa1-6, whereby BAY-876 reversed the metabolic rewiring in glycolysis, the PPP and nucleotide synthesis. In line with these findings, metabolic dysregulation is considered a key regulator of HCC, with pathways such as glycolysis, the PPP, the TCA cycle and lipid and nucleotide metabolism emerging as potential targets to inhibit tumor growth (Du et al., 2022; Satriano et al., 2019). Intercellular communication in the TME has been implicated in driving metabolic reprogramming to support tumor progression, by supplying growth factors, ECM components, nutrients and signaling molecules (Zhang et al., 2025). More specifically, HSCs have been implicated in this process by providing components such as growth factors, fatty acids and the glycolytic enzyme HK1 to aid in HCC growth (Q. Chen et al., 2022; Myojin et al., 2021; Zhang et al., 2025). Our study now identifies a specific role of HSCs in rewiring glucose metabolism in adjacent cancer cells, which we link to an enhanced growth. This illustrates the importance of HSC-HCC metabolic crosstalk in mediating metabolic alterations in HCC and underscores the importance of understanding this crosstalk to identify novel (metabolic) therapeutic targets that can be exploited to advance HCC treatment.

We further identify HIF signaling as a prominent and targetable mechanism underlying the increased proliferation and metabolic rewiring in Hepa1-6 cells. HIF signaling and its downstream target genes are activated when prolyl hydroxylase domain (PHD) proteins are inhibited, resulting in HIF-1/2α stabilization (R. Chen et al., 2024). We indeed observed increased stabilization and translocation of HIF-1α to the nucleus in co-cultured Hepa1-6 cells. In addition, interaction with mHSCs induced the expression of *Vegfa* and *Slc2a1*, two important HIF target genes (Rashid et al., 2021). Finally, suppressing HIF-1α and HIF-2α signaling by PX-478 and PT-2385, respectively, decreased proliferation of co-cultured Hepa1-6, with PT-2385 dampening glucose usage in glycolysis. This suggests that both HIF-1α and HIF-2α may contribute to the metabolic and proliferative effects of mHSC-Hepa1-6 crosstalk. Indeed, both *Hif-1α* and *Hif-2α* are generally expressed in Hepa1-6, which is in line with the involvement of both isoforms in HCC (L. Ma et al., 2014). Given that HIF-1α and HIF-2α have common targets but also non-redundant functions, isoform-specific inhibition may be partially compensated by the activity of the other and will likely have less effect than targeting all isoforms at once. The partial effect of HIF-2α inhibition on the metabolic rewiring may therefore reflect the simultaneous upregulation of both isoforms that would result in residual HIF-1α activity. Our observations are consistent with previous reports identifying HIF as an important regulator in HCC (R. Chen et al., 2024; Husain et al., 2022; Li et al., 2021; Rashid et al., 2021; Sin et al., 2023). HIF has also been associated with driving metabolic reprogramming in HCC, especially pathways involved in glucose metabolism such as glycolysis (Husain et al., 2022; J. Liu et al., 2025; Sin et al., 2023). These alterations are linked to poor prognoses, with high expression of *VEGFA* and *SLC2A1* in HCC being associated with lower patient survival rates (Lian et al., 2018).

Although hypoxia represents a *bona fide* activator of HIF signaling in cancers, HIF can also become activated in the absence of actual hypoxia, as also suggested in our co-culture system. CAFs in HCC arise predominantly from activated HSCs, which are known to secrete a broad range of cytokines that can shape tumor cell behavior (Molenaar et al., 2023; Tsuchida & Friedman, 2017). Several HSC-derived factors, including CCL5, IL-6, and TGF-β have been implicated in the activation of HIF signaling in HCC cells (H. Ma et al., 2018; H. Xu et al., 2022). However, we did not observe significant induction of CCL5, IL-6, or TGF-β signatures in our differential gene expression analysis in either co-cultured Hepa1-6 cells or mHSCs. We also did not find an upregulation of *Tgm2* in co-cultured Hepa1-6, which was reported to activate HIF signaling in HCC cells through HSC-derived IL-6 and TGF-β (H. Ma et al., 2018). This together suggests that alternative mechanisms may contribute to the HIF activation in our model. In addition to cytokines, oxidative stress represents another potential mechanism linking HSCs to HIF activation in HCC cells. ROS can stabilize HIF, contribute to metabolic reprogramming and epithelial-mesenchymal transition (EMT), and promote cancer progression (Cai et al., 2026; Z. Liu et al., 2017; Movafagh et al., 2015). We identified mHSC-derived H_2_O_2_ as a candidate crosstalk ROS that mediates HIF activation in Hepa1-6. The exact source of the mHSC-derived H_2_O_2_ remains unresolved, but may reflect a combination of decreased H_2_O_2_ scavenging and increased intracellular or extracellular ROS-generating activities, including those of cytochrome P450 enzymes, xanthine-oxidase, lipoxygenases and lysyl oxidase (LOX). Independent of HIF activation, ROS can inhibit pyruvate kinase M2 by cysteine oxidation, thus diverting glucose flux to the PPP and enhancing nucleotide synthesis and glutathione levels (Anastasiou et al., 2011). Importantly, the ability of stabilized vitamin C to attenuate the proliferation of Hepa1-6 cells in co-culture with mHSCs, phenocopying HIF inhibition, provides functional support for a link between redox imbalance and HIF-dependent HCC proliferation. Because vitamin C is a direct cofactor of PHD-dependent HIF degradation (Miles et al., 2015), it can suppress HIF signaling by both scavenging ROS and supporting HIF degradation. This is supported by previous work illustrating the potential of targeting redox signaling and scavenging excessive ROS, to promote HIF degradation and inhibition of EMT in HCC (Sun et al., 2021). Our findings do not establish whether ROS acts upstream of HIF or whether both HIF-suppressing processes are independently affected. Nevertheless, our data support a model in which HSC-derived ROS contributes to HIF activation and metabolic adaptation in HCC cells, thereby promoting their proliferative phenotype during co-culture.

As anti-angiogenic therapies targeting VEGF represent the cornerstone of current systemic HCC treatment (Cerreto et al., 2023; Selene et al., 2024), our findings provide a new perspective on the regulation of HCC angiogenesis by pointing toward an H_2_O_2_-driven HIF-VEGF signaling axis through HSC-dependent crosstalk. While HIF-mediated VEGF signaling in HCC angiogenesis is traditionally linked to tumor cell-induced hypoxia, our findings suggest that H_2_O_2_ derived from HSCs may provide an additional mechanism driving this pathway. Targeting this upstream HSC-dependent mechanism may therefore provide a more effective approach to modulating HIF-VEGF signaling than the current downstream targeting of VEGF, potentially limiting compensatory responses. This may reveal therapeutically exploitable targets, particularly through further elucidation of the metabolic mechanism underlying H_2_O_2_ production in HSCs.

We acknowledge that our study has limitations that warrants further studies. While our findings advance and strengthen the current understanding of HSC-HCC interactions and their contribution to malignant characteristics of HCC, our co-culture model also represents a simplified model using murine cell lines. Studying the HSC-HCC crosstalk in more physiologically relevant models such as human patient-derived organoid systems would further improve our understanding of HSC-driven metabolic remodeling in HCC. In addition, as we identify HSC-derived H_2_O_2_ as a promising candidate facilitating HSC-HCC crosstalk, the current findings do not yet establish a direct causal link between H_2_O_2_ and all observed downstream events. By utilizing a variety of compounds with ROS-scavenging capacities in co-culture, assessing if co-culture effects are abolished upon treatment with catalase (that specifically targets H_2_O_2_), and confirming ROS-dependent HIF activation, the signaling loop of HSC-HCC crosstalk proposed in this work could be further substantiated. Future research could also include analyses beyond polar metabolites, such as lipidomics and/or untargeted metabolomics, to provide a more comprehensive characterization of metabolic pathways involved in HSC-HCC communication and metabolic rewiring.

In conclusion, our study established a robust HSC-HCC crosstalk system to model the TME of HCC, and identified HSC-dependent HIF activation and glucose metabolism rewiring in HCC cells as a mechanism of tumor cell proliferation. We further demonstrate HSC-derived oxidative stress, with H_2_O_2_ as a candidate crosstalk ROS within the TME as a potential mechanism of pseudohypoxic HIF activation in HCC. Collectively, our findings highlight HSC-HCC crosstalk as a previously underappreciated driver of HCC progression and suggest that targeting stromal-tumor communication may offer a novel therapeutic strategy beyond direct targeting of tumor-cell-intrinsic pathways.

## Acknowledgements

The authors thank Jeroen Jansen at the Utrecht Metabolism Expertise Centre (Utrecht University) for technical support and advice during mass spectrometry experiments, and Richard Wubbolts and Esther van’T Veld of the Centre at the Centre of Cell Imaging (Utrecht University) for their invaluable assistance with immunofluorescence imaging and subsequent signal quantification. We acknowledge the Utrecht Sequencing Facility (USEQ) for providing sequencing service and RNA-seq data. USEQ is subsidized by the University Medical Center Utrecht and The Netherlands X-omics Initiative (NWO project 184.034.019).

