## Supplemental Figures for "Hepatic stellate cells drive hypoxia-inducible factor (HIF)-dependent metabolic rewiring and proliferation in hepatocellular carcinoma cells"

**Supplemental Figure S1**

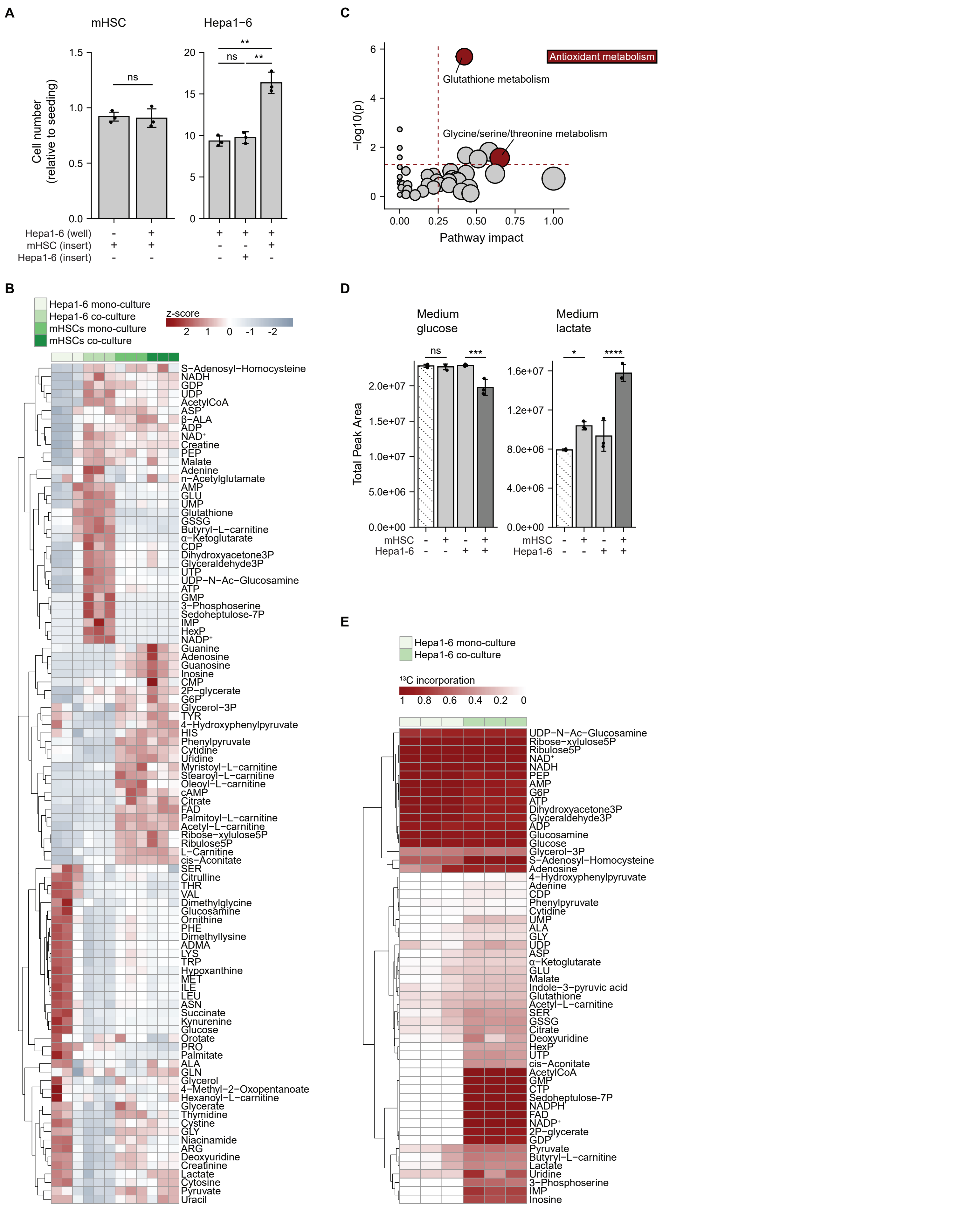


**Supplemental Figure S1.**

For panels **B-E**, metabolomics data were obtained from co- and mono-cultured mHSCs and Hepa1-6 cells after 48 hours of culturing and a 24-hour incubation with $[U-^{13}C_6]$-D-glucose (n = 3 replicates per condition). **A)** Proliferation rates of co- and mono-cultured mHSCs (left panel) and Hepa1-6 cells (right panel) (72 hours), depicted as fold change of cell number relative to the seeding density (n = 3 technical replicates per condition). A control condition showing a ‘co-culture’ condition of Hepa1-6 + Hepa1-6 is also included. Data are presented as mean ± SD. An unpaired t-test with Welch’s correction was used for statistical analysis, significance is indicated by asterisks (** = $p < 0.005$). **B)** Hierarchical clustering of all detected intracellular metabolites. Values are presented as z-scores of total metabolite levels. **C)** Pathway impact analysis of all identified intracellular metabolites, comparing here co- and mono-cultured mHSCs using MetaboAnalyst. Pathway impact and significance are depicted on the x-axis and y-axis respectively, with p-values depicted as -log10 values. Pathways highlighted in red are oxidative stress-related pathways with $p < 0.05$ and a pathway impact ≥ 0.25. **D)** Total levels of medium glucose and lactate across experimental conditions, including a medium-only control (bar with dotted pattern). Data are presented as mean ± SD. Statistical significance was assessed using a two-way ANOVA, which showed a significant interaction in co-culture with a p-value of 0.0033 and 0.0057 for glucose uptake and lactate secretion respectively. This was followed by a Šídák’s multiple comparisons test, from which its significance is indicated in the plot, illustrated by asterisks (* = adjusted $p < 0.05$, *** = adjusted $p < 0.001$, **** = adjusted $p < 0.0001$). **E)** Hierarchically clustered heatmap showing the fraction of $^{13}C$ incorporation across intracellular metabolites, comparing here co- and mono-cultured Hepa1-6. Metabolites were filtered for at least 5% of label incorporation in any sample.

### Supplemental Figure S2

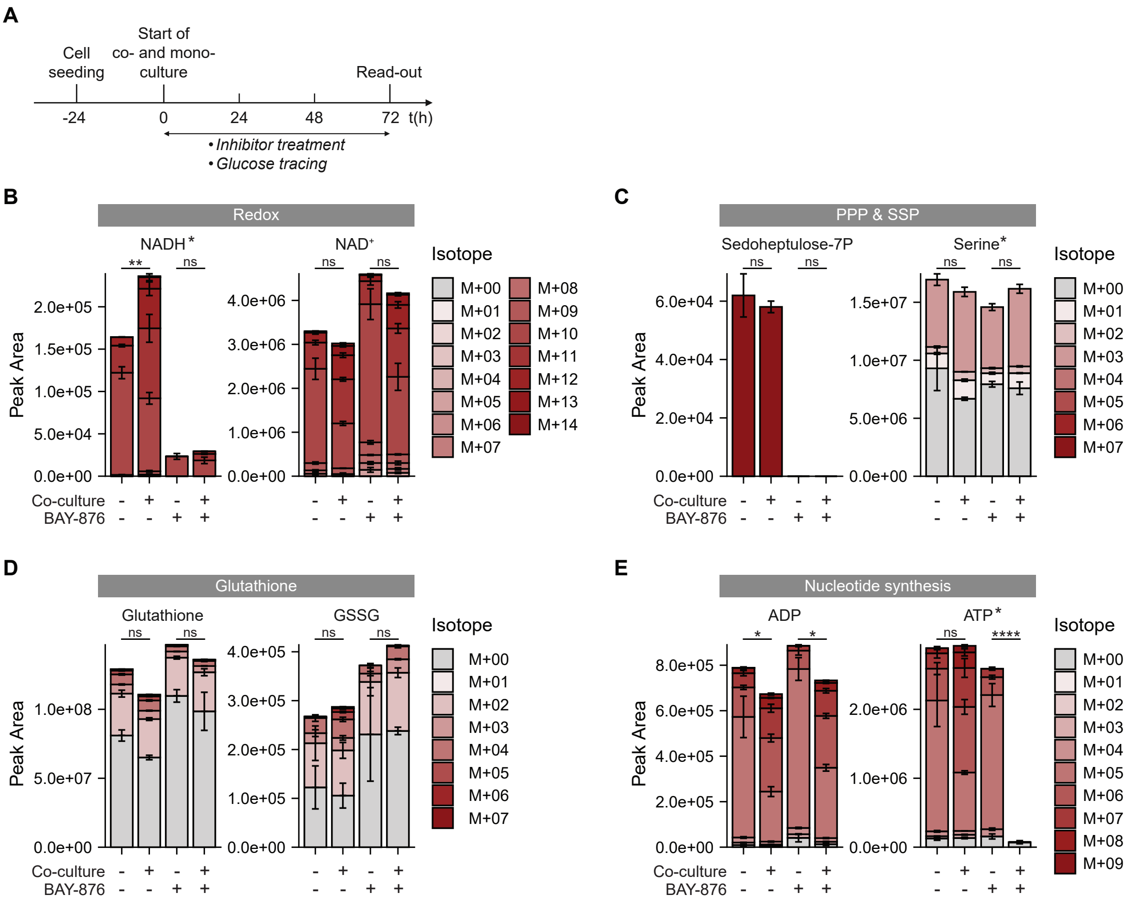


**Supplemental Figure S2.**
**A)** Timeline of the Transwell co-culture model. Hepa1-6 cells and mHSCs were seeded 24 hours prior to starting the co- and mono-cultures. Subsequently, cells were cultured for an additional 72 hours, at which point read-outs were performed. Throughout the 72-hour culture period, cells were subjected to inhibitor treatment and incubation with [U-$^{13}C_6$]-D-glucose for isotope tracing. **B, C, D, E)** Levels and isotopologue distribution patterns of intracellular redox metabolites **(B)** PPP and SPP metabolites **(C)**, (oxidized) glutathione **(D)**, and purines **(E)**, in co- and mono-cultured Hepa1-6 cells after this 72-hour incubation with [U-$^{13}C_6$]-D-glucose in the presence or absence of BAY-876 (5 nM) (n = 3 replicates per condition). Data are presented as mean ± SD. Statistical significance was assessed using a two-way ANOVA, which showed a significant interaction between culture (mono-/co-culture) and inhibitor (no inhibitor/BAY-876) conditions for metabolites with an asterisk, followed by Šídák-adjusted post hoc multiple comparisons based on estimated marginal means. Significance shown in the figure corresponds to these post hoc comparisons, illustrated by asterisks (* = p < 0.05, ** = p < 0.01, **** = p < 0.0001).

**Supplemental Figure S3**

**A**

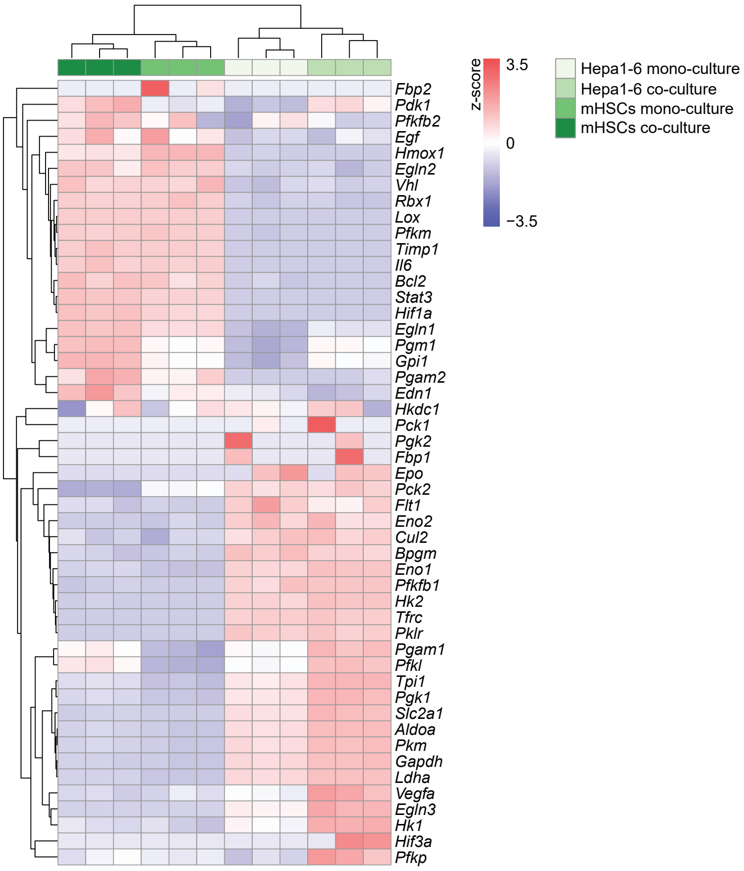


**Supplemental Figure S3.**
RNA sequencing data of Hepa1-6 and mHSC cells, obtained after 72 hours of mono- or co-culture. Raw readcounts were normalized using The Utrecht University ATWiCS ShinyApp. **A)** Hierarchically clustered heatmap showing z-score-normalized gene expression of genes related to HIF signaling for co- and mono-cultured Hepa1-6 and mHSCs (n = 3 replicates per condition).

### Supplemental Figure S4

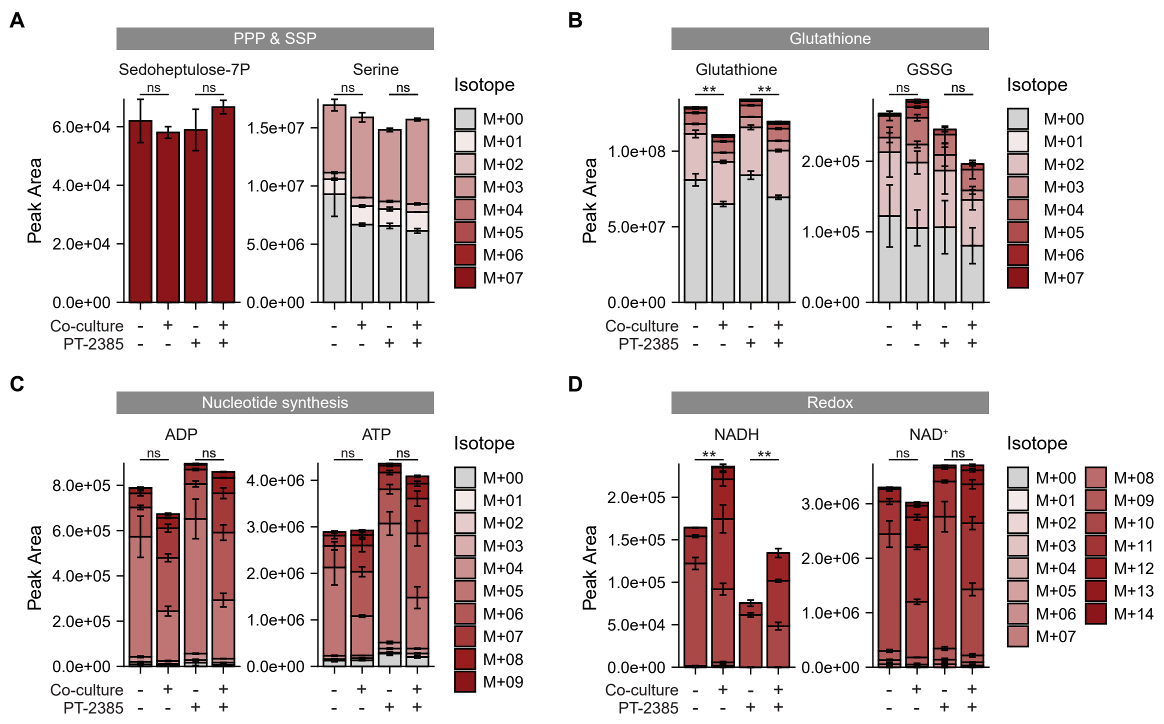


**Supplemental Figure S4.**
**A, B, C, D)** Levels and isotopologue distribution patterns of intracellular PPP and SPP metabolites **(A)**, (oxidized) glutathione **(B)**, purines **(C)** and redox metabolites **(D)**, in co- and mono-cultured Hepa1-6 cells after a 72-hour incubation with [U-$^{13}C_6$]-D-glucose in the presence or absence of PT-2385 (10 µM) (n = 3 replicates per condition). NB: PT-2385 and BAY-876 conditions were combined in one metabolomics experiment. Control (untreated) co- and mono-cultured Hepa1-6 cells are therefore identical to those presented in Fig. 2 and S2. Data are presented as mean ± SD. Statistical significance was assessed using a two-way ANOVA, which showed a significant interaction between culture (mono-/co-culture) and inhibitor (no inhibitor/ PT-2385) conditions for metabolites with an asterisk, followed by Šídák-adjusted post hoc multiple comparisons based on estimated marginal means. Significance shown in the figure corresponds to these post hoc comparisons, illustrated by asterisks (** = $p < 0.01$).

### Supplemental Figure S5

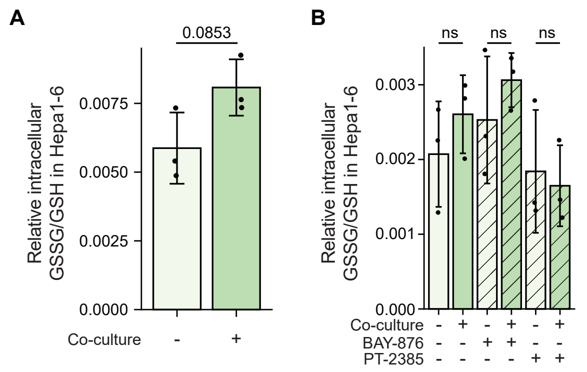


**Supplemental Figure S5.**
**A)** Relative GSSG/GSH ratio in co- and mono-cultured Hepa1-6 cells after 48 hours of culturing and a 24-hour incubation with [U-$^{13}C_6$]-D-glucose (n = 3 replicates per condition). Values were calculated using total peak areas of GSH and GSSG as assessed by metabolomics (data from the same experiment are also presented in Fig. 1 and S1). Data are presented as mean ± SD. An unpaired t-test with Welch's correction was used for statistical analysis. **B)** Relative GSSG/GSH ratio in co- and mono-cultured Hepa1-6 cells after a 72-hour incubation with [U-$^{13}C_6$]-D-glucose in the presence or absence of BAY-876 (5 nM) or PT-2385 (10 µM) (n = 3 replicates per condition). Values were calculated using total peak areas of GSH and GSSG as assessed by metabolomics (data from the same experiment are also presented in Fig. 2, 4, S2 and S4). Data are presented as mean ± SD. Unpaired t-tests with Welch's correction were used for statistical analysis.
